# Rapamycin treatment during development extends lifespan and healthspan

**DOI:** 10.1101/2022.02.18.481092

**Authors:** Anastasia V. Shindyapina, Yongmin Cho, Alaattin Kaya, Alexander Tyshkovskiy, José P. Castro, Juozas Gordevicius, Jesse R. Poganik, Steve Horvath, Leonid Peshkin, Vadim N. Gladyshev

## Abstract

The possibility that pace of development is tightly connected to aging is supported by the fact that the onset of reproduction is associated with lifespan and that many longevity interventions target growth and development. However, it has been unknown whether targeting development with pharmacological intervention can lead to a longer lifespan. To test this possibility, we subjected genetically diverse UMHET3 mice to the mTOR inhibitor rapamycin for the first 45 days of life and followed them up until death. Treated mice grew slower, with most of the deceleration occurring in the first week, and remained smaller for their entire lives. Their reproductive age was delayed but without affecting offspring numbers. The treatment was sufficient to extend the median lifespan by 10%, with the most effect in males, and to preserve better health as measured by frailty index, gait speed, and glucose and insulin tolerance tests. Mechanistically, the liver transcriptome of treated mice was younger at the completion of treatment and stayed younger into the old ages in males. Rapamycin initially reduced DNA methylation age of livers, however, that effect was lost with aging. Analogous to mice, rapamycin exposure only during development robustly extended the lifespan of *Daphnia magna* as well as reduced their body size, suggesting evolutionary conserved mechanisms of this early life effect. Overall, the results demonstrate that short-term rapamycin treatment during early life is a novel longevity intervention that establishes causality between pace of development and longevity in evolutionary distant organisms.

## Introduction

Evolutionary history of mammals is a prominent example of how lifespan may be changed over 100-fold. Cross-species analyses revealed a strong positive association of species’ maximum lifespan with time to maturity and a positive association with body mass (1, 2). On the contrary, smaller animals within species tend to live longer or be protected from age-related diseases as evidenced by studies of dogs and humans (3–6). These associations could be explained by the importance of growth rate for longevity (1, 2). Indeed, knockouts of the growth hormone pathway or genes involved in insulin/IGF-1 signaling (IIS) lead to extended lifespan of organisms from worms to mice (7–13), and this is further associated with decreased body size. Moreover, wild type mice preselected for slower growth live longer than their fast-growing siblings (14). Likewise, the growth rate of longer-lived species of mammals is lower compared to those with shorter lifespan (15). These findings suggest a causal link between the organismal pace of growth, body size and longevity.

Some indirect evidence supports the causal relationship between inhibition of growth signaling and longevity if targeted during development. For example, growth hormone knockout mice and mice lacking GH production live up to 50% longer than their wild type siblings (10, 12). However, their longevity was diminished if they were treated with growth hormone during early postnatal development (16, 17). At the same time, growth hormone knockout induced at adult age had limited to no effects on longevity (18). However, there have been no experiments where growth pathways are directly inhibited only during development and the longevity outcomes measured.

Rapamycin is a well-characterized mTOR inhibitor and is among the most validated and potent pharmaceutical interventions that extend lifespan in mice. Rapamycin can extend lifespan if given in adulthood (19, 20) or later in life (21, 22) in various mouse strains, including genetically diverse UMHET3 mice (a cross of four inbred strains). Interestingly, rapamycin failed to extend the lifespan of growth hormone receptor knockout mice (20). Furthermore, early life rapamycin treatment was previously shown to suppress growth of mice (23). Thus rapamycin is a perfect candidate to test how targeting growth only early in life can affect lifespan, and we employed it in our study, examining its effects on longevity, healthspan, biological age and gene expression.

## Results

### Rapamycin treatment during mouse development attenuates growth and onset of reproduction

To evaluate the effect of rapamycin on longevity, we set up a cohort of 130 newborn UMHET3 mice for a lifespan experiment and a separate cohort of 40 mice for healthspan analyses (Fig. 1A). Dams with newborns were fed chow containing 42 ppm of eudragit-encapsulated rapamycin or the corresponding amount of encapsulating material as a control. Pups were separated from dams at 20-22 days of age and continued on the same rapamycin diet until 45 days of age (Fig. 1A). These animals were then transferred to a regular chow diet and followed for the rest of their lives.

**Figure 1.**
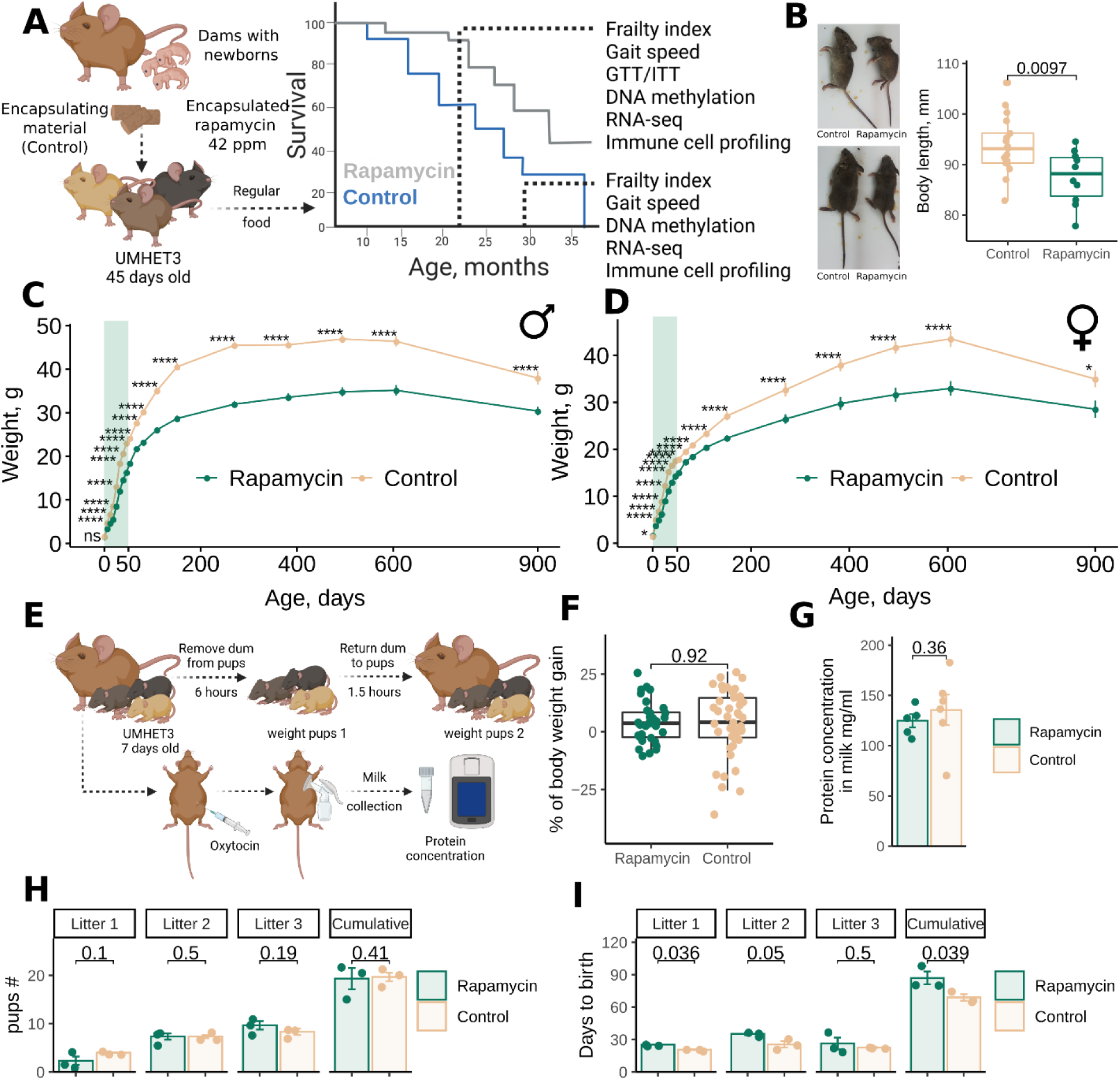
Rapamycin inhibits growth and delays reproduction of genetically heterogeneous mice. (A) Experimental design for evaluating the effect of early life rapamycin treatment on mouse longevity and health. (B) Representative images of 2-month-old UMHET3 mice subjected to eudragit-encapsulated rapamycin or control (Eudragit-containing) diets during the first 45 days of their lives. (C)-(D) Body weight change of male (C) and female (D) mice subjected to rapamycin (n=30 per sex) or control (n=34-39 per sex) diets from 0 to 45 days of age. The shaded area shows the period of rapamycin treatment. The data are mean ± SE. (E) Experimental design for evaluating the effects of early life rapamycin treatment on pups’ nutrition. (F) Percentage of 7-day old pups’ body weight gain calculated as weight of pup after it consumed milk divided by the mean weight of the group after 6-hour starving period and exactly before pups were returned to dams and multiplied by 100. Mean weight was used for normalization within each group because of the baseline difference in weight between rapamycin and control pups. (G) Milk protein concentration in dams subjected to rapamycin or control diets starting from the day they give birth. (H) Number of pups born to breeders subjected to rapamycin or control diets during development. (I) Time between pregnancies for rapamycin and control groups of breeders.

Rapamycin treatment in early life (EL) resulted in smaller mice (Fig. 1B), and the reduced weight was preserved across the lifespan in both males (Fig. 1C) and females (Fig. 1D). Rapamycin showed an effect within days. Small body size was seen upon completion of treatment, with males being affected more than females. Accordingly, the weight of organs (brain, liver, kidney, spleen) was also reduced in the treatment group (Fig. S1A).

Because rapamycin was initially delivered to the pups through the milk, we examined whether rapamycin directly inhibited the growth of animals or affected nutrient consumption. Thus, we tested if pups consumed the same amount of milk and if total protein of the milk was affected in the treated animals (Fig. 1F). We focused on protein levels because rapamycin inhibits protein synthesis (24, 25). We found no difference in milk consumption between treated and control groups (Fig. 1F); each group gained the same percentage of weight after being separated from dams and returned for refeeding. We further constructed a custom milking pump and collected milk from the dams (Fig. S2). The total protein content of the milk was not different between treated and control dams (Fig. 1G). We conclude that the observed effect of growth patterns is the direct consequence of rapamycin treatment rather than nutritional deficits.

Growth is usually coupled with time to sexual maturity. To test whether EL rapamycin treatment not only attenuated body growth but also the age at sexual maturity, we set up breeding pairs of treated and untreated age-matched UMHET3 mice (46-49-day old females and 47-day old males) and recorded the dates when the females gave birth to first, second, and third litters as well as the number of pups born. Treated and untreated mice had the same litter size (Fig. 1H). However, treated breeders had their first litters on average 4.6 days later than untreated breeders (p=0.036). The period between the first and second litters was also longer in rapamycin-treated mice (now by 9.6 days, p = 0.05)), and the period between the second and third litters was not statistically significantly (p=0.5) longer in rapamycin-treated breeders (Fig. 1I).

### EL rapamycin treatment during development extends lifespan and healthspan of mice

We found that EL rapamycin treatment extended the median lifespan of UMHET3 mice by 10% (Fig. 2A, p = 0.036). This effect was primarily due to the effect of treatment in males (Fig. 2B, p = 0.0064, median lifespan extension by 11.8%), whereas females did not live longer (Fig. 2C, p = 0.55).

**Figure 2.**
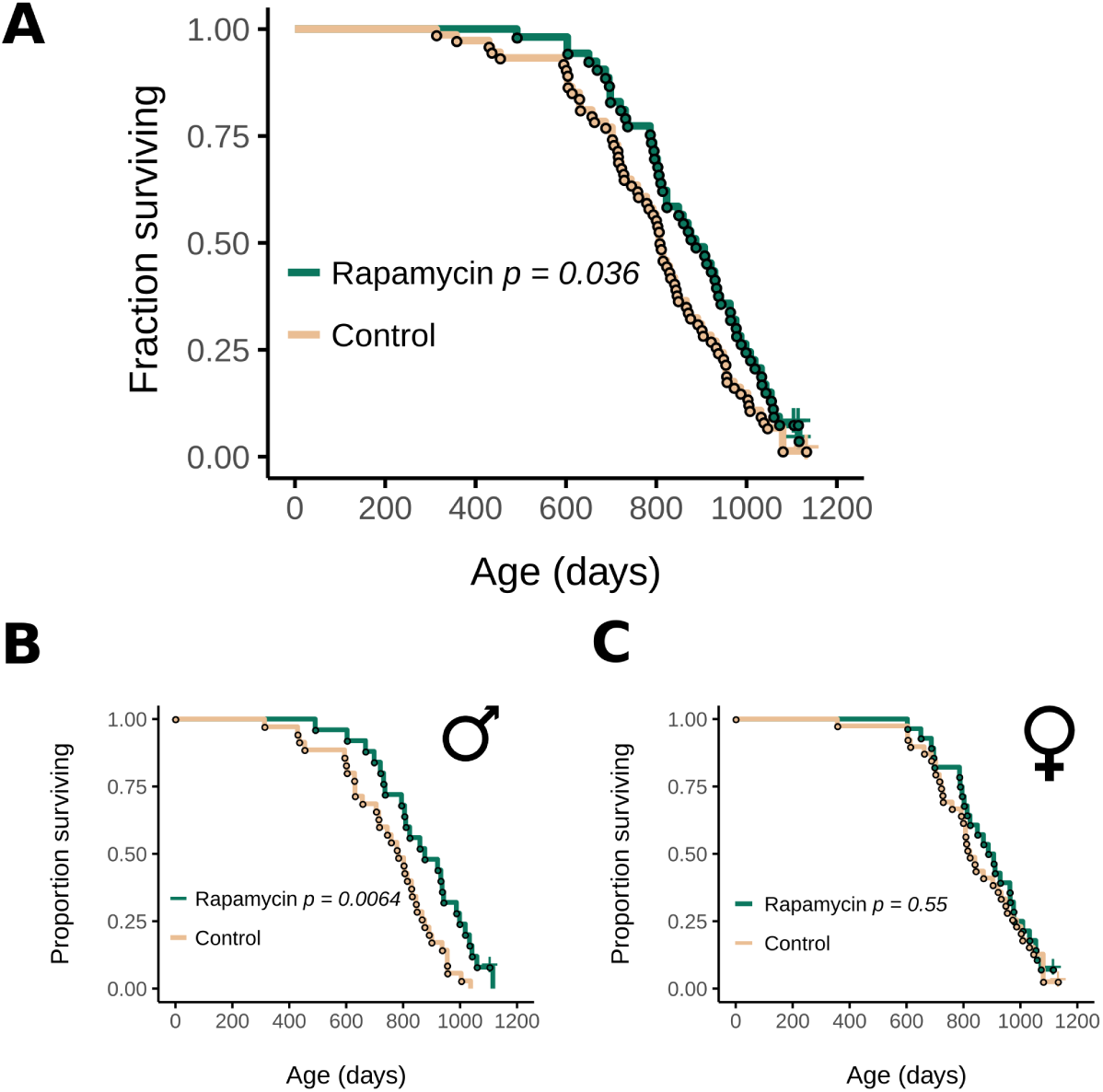
Rapamycin treatment during development extends the lifespan of UMHET3 mice. (A) Survival curve of treated and untreated UMHET3 mice, sexes combined. (B)-(C) Survival curves of male (B) and (C) female UMHET3 mice that were subjected to rapamycin or control diets from birth until 45 days of age. P-values were calculated with the log-rank test.

To understand if healthspan was also improved in the EL treated mice later in their lives, we measured frailty index (26–28), gait speed and glucose and insulin tolerance at the age of 20-29 months. Frailty index is a 31-parameter estimate of overall frailty of mice based on non-invasive assessment of mouse age-associated features (26–28). EL rapamycin treatment made mice less frail than the control group (Ancova adjusted for sex and age, p-value = 0.007, Fig. 3A). The effect was slightly more significant for males than for females (Fig. 3B), in agreement with the survival data. We further applied a frailty index clock (26) that predicts the survival of mice based on frailty score features and takes mouse age into account, and found that mice treated with EL rapamycin had a longer predicted survival time (Ancova adjusted for sex, p-value = 0.015, Fig. 3C). We conclude that EL rapamycin treatment delays the onset of frailty phenotypes, including features that are associated with mouse survival. Feature analyses revealed that the forelimb grip strength, gait disorders, body condition score and distended abdomen were improved in the treatment group (Fig. S3A).

**Figure 3.**
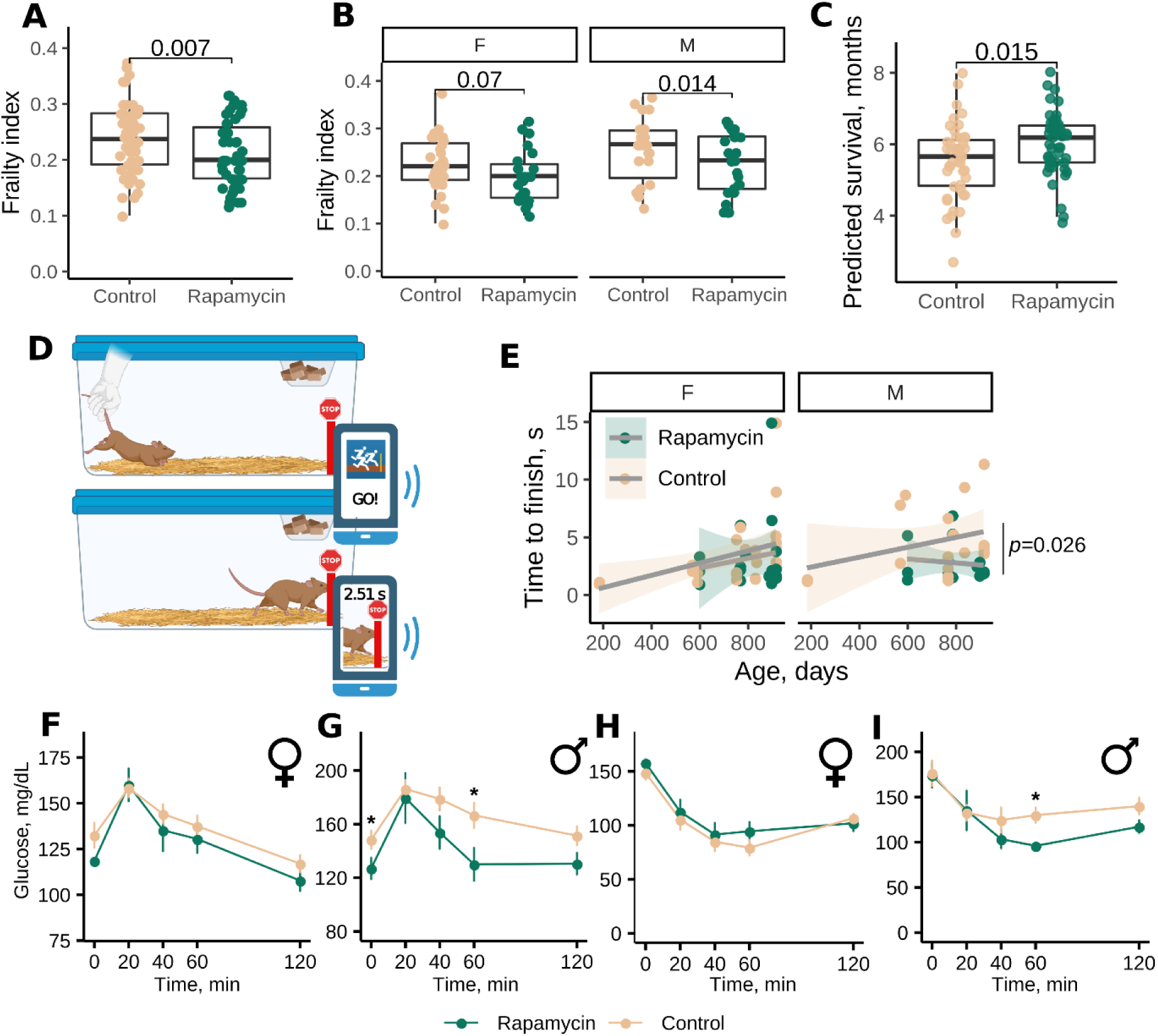
Rapamycin treatment during development extends the healthspan of mice. (A) Frailty index score of 20-29-month-old UMHET3 mice subjected to diets containing encapsulated rapamycin or encapsulating material from birth until 45 days of age. P-value was calculated with ANCOVA test and adjusted for sex and age. (B) Frailty index score of 20-29-month-old male (M) and female (F) UMHET3 mice subjected to rapamycin or control diets from birth until 45 days of age. P-value was calculated with ANCOVA test and adjusted for age. (C) Predicted survival (in months) based on frailty index calculated using the AFRAID clock. (D) Scheme of gait speed measurement in mice. (E) Median time mice spent in seconds to reach the finish line. P-value was calculated with ANCOVA test and adjusted for age. (F)-(G) Glucose tolerance test results for females (F) and (G) males (n=4-6 per group per sex). Insulin tolerance test result for (H) females and (I) males (n=4-6 per group per sex). P-values were calculated with a one-tailed Student t-test.

We further developed and applied a new test to assess mobility fitness in mice that does not require specific equipment and can be conducted by using a cell phone and a common app (Fig. 3D). Such a test can be performed at a house facility and at near zero cost. In this test, mice are placed at one end of the cage and allowed to run voluntarily to the other end of the cage (Fig. 3D). The time to finish is determined using a photo-finish app. We found that mice ran significantly slower with age (Fig. S3B). EL rapamycin attenuated this phenotype in aged males (Fig. 3E), suggesting that mobility fitness is better preserved in these animals as a result of the intervention.

Additionally, we analyzed if our EL intervention affects the metabolic health of the old animals. Glucose and insulin tolerance tests revealed some sex-specific improvements of glucose and insulin tolerance in treated males of older ages (Fig. 3F-I), consistent with sex specificity of the treament on lifespan.

Finally, we analyzed immune cell composition of the spleen and bone marrow of treated and control groups of animals 20-29 months of age. We previously established new age-related phenotypes of the immune system that are predictive of mouse lifespan and lymphoma incidence (increased B-cell size, clonal B cells, age-associated B cells and myeloid bias) (29). Analysis of control mice revealed that UMHET3 mice accumulate the same age-related phenotypes as C57BL/6 mice (B-cell size increase, myeloid skewing, accumulation of age-associated clonal B cells (ACBC) and age-associated B cells (ABC)). We found that these phenotypes (Fig. S4A,B,D,E), as well as percentage (Fig. S4C) and cell size (Fig. S4F) of hematopoietic stem cells (HSC) were not affected in the treated mice in comparison to controls. Finally, necropsy analyses of aged mice that died of natural causes revealed no difference in the incidence of common mouse age-related pathologies (Table S1), indicating that the longevity effect of the EL rapamycin treatment cannot be attributed to the prevention of a particular age-related disease.

### EL rapamycin treatment induces pro-longevity gene expression changes that are preserved with age in a sex-specific manner

To gain insights into molecular mechanisms of the EL rapamycin regimen, we analyzed gene expression changes in response to the treatment in the livers of young (2 mo), middle-aged (20-22 mo) and old (28-29 mo) mice. We first found that the number of differentially expressed genes triggered by treatment was reduced with age (Fig. S5A). Principal component analysis revealed that first component explains sex and second component explains age variation of animals (Fig. 4A). We then compared transcriptome changes in response to EL rapamycin to those induced by known longevity interventions as well as those induced by aging (i.e. compared gene expression signatures) (30). In both males and females, EL rapamycin triggered transcriptome changes in the young liver that were opposite to age-related changes in this organ and were similar to those induced by longevity interventions (lifelong rapamycin and growth hormone deficiency) (Fig. 4B). In males, a negative association with aging signatures persisted throughout life; however, in females the effect of treatment vanished with time (Fig. 4B). Remarkably, a positive association of EL rapamycin effect with the effect of lifelong rapamycin treatment was lost with age in both males and females; it was strongest in young animals days after the treatment was stopped (Fig. S5D). This indicates that the effect of rapamycin on the liver transcriptome fades away over time once the treatment stops. However, other anti-aging and pro-longevity effects on the transcriptomes were preserved in treated male mice. This was supported by the negative association with the liver aging signature and positive association with the gene expression biomarkers of mice with higher median and maximum lifespan (Median and Max lifespan signatures) (Fig. 4B). Interestingly, the sex-specific lifespan and healthspan effects of the treatment were reflected in hepatic gene expression changes. In particular, males that benefited the most from the treatment retained the negative association with aging signatures across ages. Treated females, on the other hand, acquired a positive association with aging signatures at older ages.

**Figure 4.**
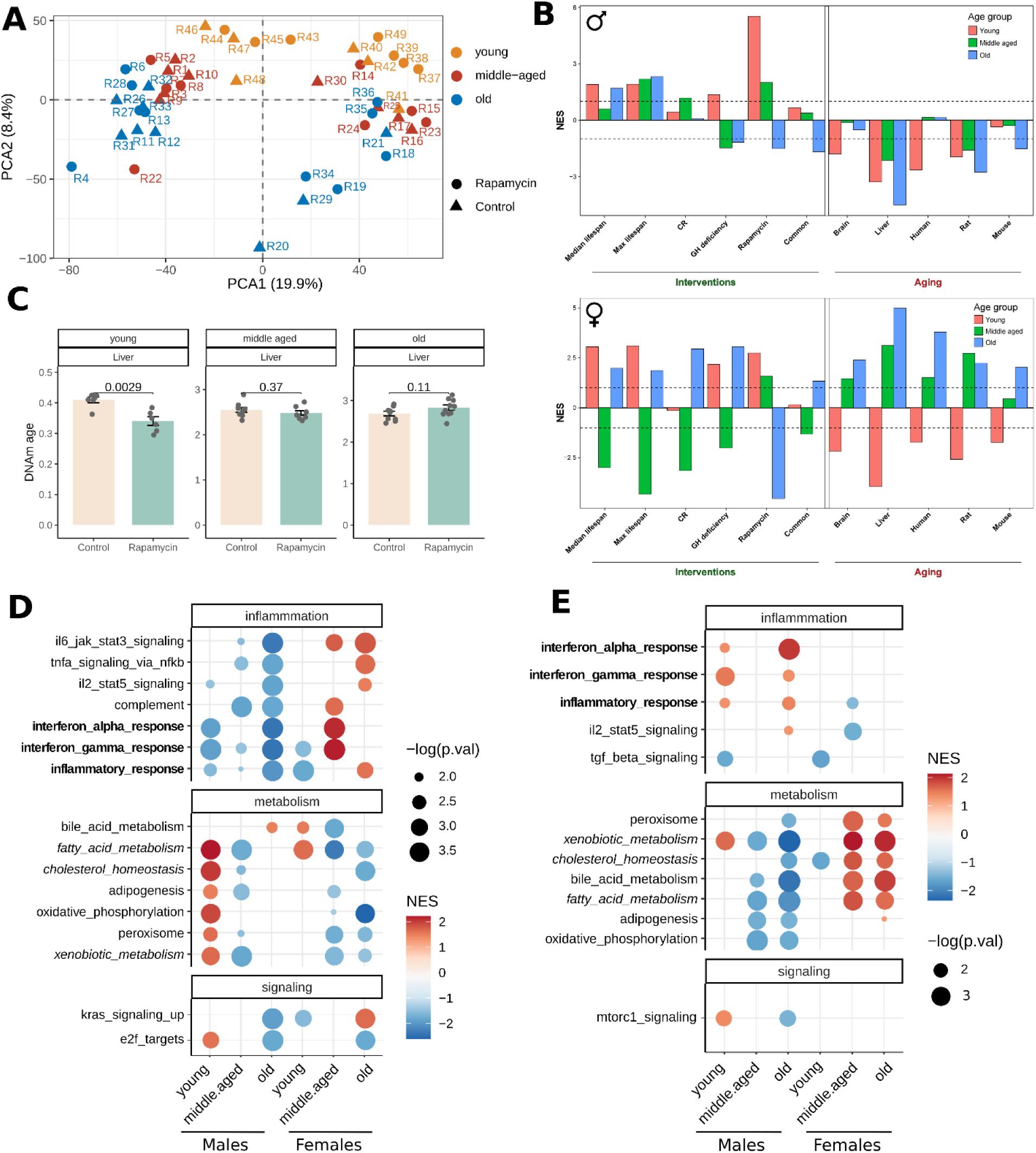
Rapamycin treatment during development attenuates aging of the transcriptome and DNA methylation of young animals, with males preserving younger transcriptomes across lifespan. (A) PCA plots of gene expression changes measured by RNA-seq. Points are shaped according to diet, and colored according to age group. All genes are used for plotting. (B) Association between gene expression changes induced by EL rapamycin and measured in young (red), middle-aged (green) and old (blue) animals, and signatures of aging and lifespan extension in males (upper plot) and females (lower plot). The intervention signatures include gene signatures of individual interventions (caloric restriction (CR), growth hormone (GH) deficiency and rapamycin), common gene expression signatures across different interventions (common) and signatures associated with the effect of interventions on maximum or median lifespan (max and median lifespan). Aging signatures include age-related gene expression changes across different tissues of humans, mice and rats (human, mouse, rat) as well as liver- and brain-specific changes (liver, brain). (C) DNA methylation age of livers estimated by the Liver epigenetic aging clock in young, middle-aged and old treated and untreated animals. P-values are calculated with a two-tailed Student T-test. (D) GSEA for hallmark pathways from msigdb database using ranked differentially expressed and (E) differentially methylated genes in treated animals. Only significant enrichments (q-values<0.1) for at least two conditions are shown. Dots are colored according to normalized enrichment score (NES) and sized according to -log10 of p-value.

Pathway gene set enrichment analysis based on the MSigDB database revealed that young animals upregulated metabolic pathways and downregulated inflammation pathways in response to the treatment in both sexes (Fig. 5D). These changes were consistent with other longevity interventions, including rapamycin at adult age, caloric restriction, growth hormone deficiency and heterochronic parabiosis (Fig. S5B-C), and were opposite to age-related changes. Consistent with the lifespan and healthspan benefits, we observed sex-specific effects on the transcriptomes. First, unlike females, treated males preserved the down-regulation of inflammation pathways with age (Fig. 4D). Second, young males upregulated metabolic pathways more robustly than young females (Fig. 4D).

**Figure 5.**
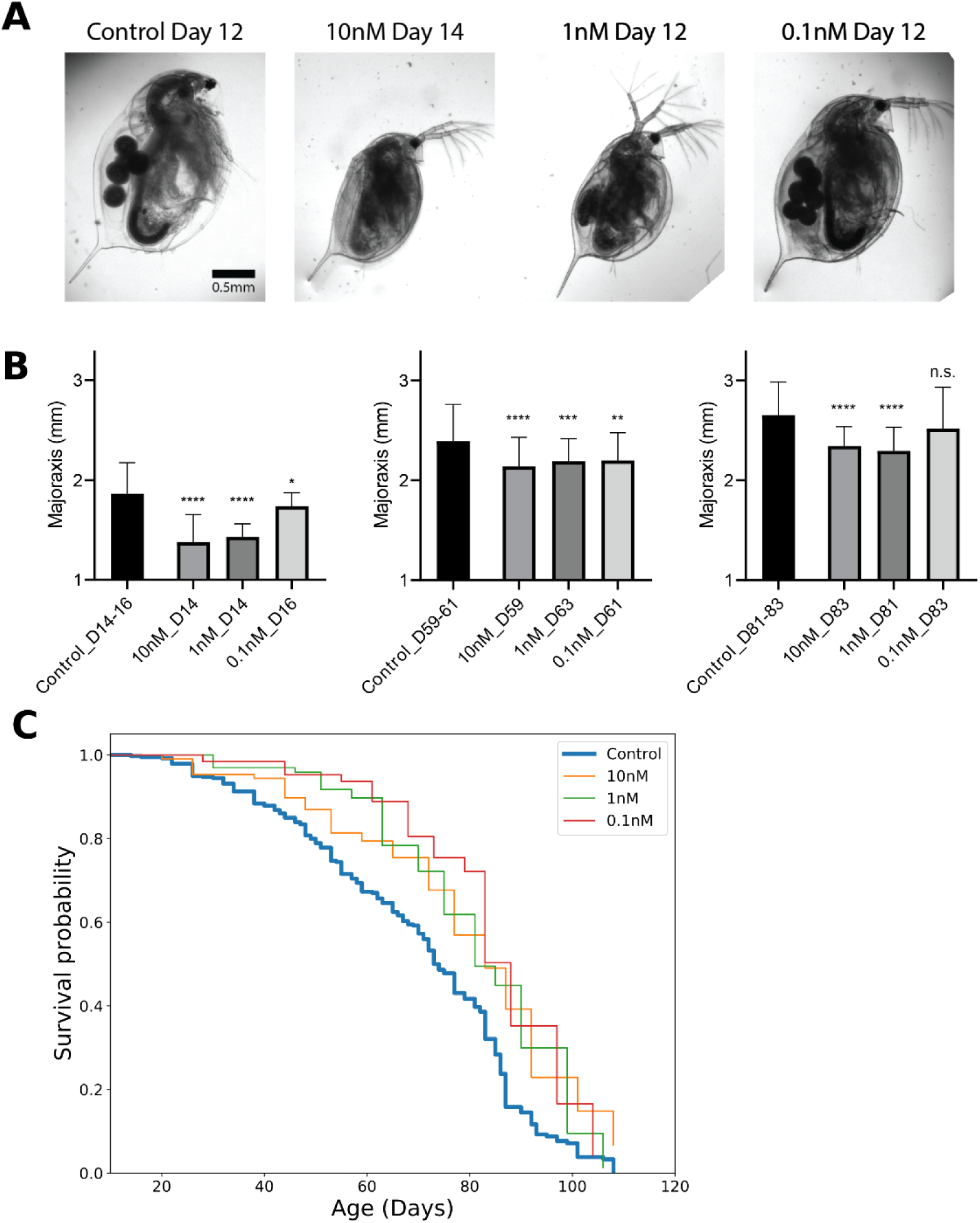
Rapamycin treatment during development extends lifespan and reduces the body size of *Daphnia magna*. (A) Representative images of *Daphnia magna* at 12-14 days following rapamycin exposure. Also shown is a control animal at the same chronological age. (B) Daphnia body size measurement at days 14-16, days 59-63, and days 81-83. p-values are calculated using two-sided Student T-test.. The data are mean ± SD. Kruskal-Wallis test (****: p<0.0001, ***: p<0.001, **: p<0.01, *: p<0.05). (C) Survival curves for control and three different concentrations of rapamycin (Control: n = 379, 0.01 nM: n = 107, 1 nM: n = 97, 0.1 nM: n = 63).

Overall, we found that the EL rapamycin treatment initially triggered gene expression changes that were similar to potent longevity interventions and were opposite to age-related changes, with males better preserving these effects across lifespan. However, females lost negative association with aging signatures at older ages, consistent with the sex-specific effects of the EL rapamycin treatment on lifespan and healthspan. Moreover, there was no correlation between gene expression ranks of old males and females (spearman correlation coefficient, *r*=-0.02), further underlying the sex-specificity of impact of EL rapamycin on gene expression into older ages of animals (Fig. S5E).

### EL rapamycin perturbes global DNA methylation patterns in mouse liver

To test a potential role of the epigenome on the long-lasting effects of EL rapamycin, we performed analyses of DNA methylation (DNAm) of the livers that were analyzed with RNA-seq. The treatment significantly affected 83 CpG sites (FDR < 0.1) independent of age and sex of the animals (Table S2). We further performed GO and KEGG enrichment of genes that carry differentially methylated CpG sites and are differentially expressed. These genes in young animals were significantly enriched in steroid, thyroid and hormone biosynthesis (Fig. S6). This finding suggests that EL rapamycin regulates the expression of genes involved in hormone biosynthesis by altering their DNAm patterns. This is consistent with the changes in body size and sex-specific effects of the treatment.

We then estimated the DNAm age of the livers using pan-tissue and liver-specific epigenetic aging mouse clocks. Interestingly, the EL rapamycin treatment initially reduced epigenetic age of the livers of young animals in both sexes but this effect was lost with age (Fig. 4C).

Enrichment of differentially methylated regions of young, middle-aged and old treated males and females revealed a limited number of enriched pathways that changed following the treatment and were also preserved across lifespan in both sexes (Fig. 4E). Moreover, correlation of CpG ranks among ages and sexes was relatively high only between young males and young females (spearman correlation coefficient, *r*=0.12), and between old males and middle-aged males (*r*=0.15) (Fig. S5F).

### Rapamycin treatment during development extends the lifespan of *Daphnia magna*

To understand if the EL rapamycin treatment operates through evolutionarily conserved longevity mechanisms, we subjected the model invertebrate *Daphnia magna* to rapamycin only during postnatal development. We chose a window from day 3-4 to day 11-12 as most of the animals begin to produce offspring by day 12 (31).

Like mice, EL rapamycin-treated Daphnia were smaller in size, and the effect was dose-dependent (Fig. 5A-B). While animals treated with 1 and 10 nM rapamycin stayed smaller throughout their lifespan, *Daphnia* treated with a 0.1 nM rapamycin caught up in size as they aged and by the age of 83 days were insignificantly different from controls (Fig. 5B, right panel). Despite catching up in size, these animals still lived longer suggesting that staying small throughout lifespan is not necessary for the longevity effect of the treatment.

Concentrations above 10 uM were found to be toxic and were not studied further. We did not observe a significant effect of 0.1 μM rapamycin on lifespan (*χ*2 = 1.505 and p-value = 0.220). We then examined the effect of three lower doses of rapamycin (10 nM, 1 nM, and 0.1 nM) on lifespan and found a significant increase in lifespan for all three conditions (control vs 10 nM: *χ*2 = 22.72 and p-value <0.0001, Control vs 1 nM: *χ*2 = 16.23 and p-value < 0.0001 and Control vs 0.1 nM: *χ*2 = 18.67 and p-value < 0.0001 in log-rank test) (Fig. 5C).

## Discussion

In this study, we demonstrated that longevity can be achieved by EL rapamycin treatment which is associated with the delay of growth and development in evolutionarily distant model organisms: mice and the invertebrate *Daphnia*. Many longevity interventions that target growth pathways and body size are inversely associated with longevity within species, and our findings provide the first causative evidence that slowing mTOR-mediated growth results in longer lifespan.

There is much evidence that early life exposures may affect aging and lifespan in evolutionarily distant organisms. For example, mouse pups that get less nutrition in the crowded litter model are longer lived, though the mechanisms involved are unknown (32). There are also some extreme cases of longevity achieved by changing the diet during development of insects. Bees fed royal jelly during development become queens and live more than twice as long as untreated worker bees (33, 34). It was also reported that natural increase in reactive oxygen species during early development extends lifespan of *C. elegans* by acting on H3K4me3 marks (35). However, none of the previous studies involved pharmacological interventions, which makes interpretation of underlying mechanisms difficult. The fact that EL exposure to rapamycin, an mTOR inhibitor, during postnatal development increases the lifespan and inhibits the growth of two evolutionarily distant organisms supports the idea that mTOR is a crucial gene for organismal growth and that it mediates the relationship between pace of growth and longevity. However, rapamycin has off-targets that also impact growth and cell division, for example, STAT3 and c-MYC in human cells (36). Future studies may test if mTOR-deficient mice respond to the EL rapamycin regimen similarly to wild type animals.

The ways early life exposures translate into longevity phenotypes are unknown. One possibility is the epigenetic memory of the treated tissues through chromatin structure and DNA methylation. In our analysis, few changes in the liver DNA methylation were retained by the treated animals all the way into old age. However, it is possible that the EL rapamycin treatment has a long-lasting impact on other epigenetic marks, for example, those that affect chromatin structure, as was recently shown for the mouse intestine (37). Alternatively, tissues other than the liver (analyzed here) may be involved in the epigenetic memory of the EL rapamycin treatment.

It is also possible that a boost of longevity mechanisms during early life alone is sufficient to attenuate aging phenotypes later in life. Both gene expression signatures and DNAm clocks revealed the strongest longevity effects of EL rapamycin in young animals, whereas these effects were mostly lost later in life. Driver somatic mutations may occur at a young age and evolve into a full blown cancer only later in life (38). Early-life treatments can prevent or delay the appearance of somatic mutations, leading to delayed age-related clonal expansion and cancer. Another explanation is an improved proteostasis. A nuclear pore complex and histones are examples of proteins that are synthesized early in life and preserved throughout organismal lifespan in non-replicative tissues (39, 40). At the same time, there is a recent finding that rapamycin improves translation precision *in vitro* (41). Rapamycin treatment may reduce the number of translation errors in long-lived proteins and improve their cumulative function throughout lifespan, even if given briefly at a young age. Future studies may test if EL rapamycin does so *in vivo*.

Another significant observation of this study was the sex-specific effect of EL rapamycin treatment in mice. Both lifespan and healthspan improvements were significant in males, while there was only a trend towards improvement in female mice. This finding is consistent with previous lifespan studies using growth manipulation. Unlike males, female mice benefited from conditional knockout of the growth hormone receptor during adulthood (18). This demonstrates that the benefits of growth hormone receptor knockout for female lifespan is a combination of early-life and late-life effects, while only early-life exposure matters for the male lifespan. Similarly, growth hormone deficient males respond more strongly to early-life growth hormone therapy than females (16). One explanation for these effects could be that females are naturally smaller and thus growth retardation will have a smaller impact on their phenotypes. Indeed, we found that EL rapamycin impacted growth of male mice more than females. Another explanation could be a non-trivial relationship between molecular mechanisms of growth control, sex and longevity. In this study, we found that treated females reduce inflammation pathways at first but overactivate them later in life as compared to controls. This indicates an active gain of detrimental functions at the old age alongside the loss of beneficial effects from the treatment. Lastly, in the mouse experiment, only one dosage of rapamycin was tested. However, our Daphnia experiments revealed that lower dosages can be as beneficial as the higher ones, while the highest rapamycin concentrations were detrimental. Future studies may explore the effects of various dosages of EL rapamycin on mouse lifespan.

### Experimental Procedures

#### Animals

CByB6F1/J females and C3D2F1/J males mice were purchased from the Jackson Laboratory. Breeder pairs were set up and monitored daily. On the day of birth, males were removed from the cages, and the dams were given chow containing encapsulated rapamycin (42 ppm of active compound) or encapsulating material Eudragit at the same concentration (Rapamycin Holdings) in 5053 diet (TestDiet) and were fed *ad libitum* until 45 days of age. Then, all mice were subjected to a regular 5053 diet (TestDiet) and followed until death or moribund state. Mice that died as a result of fighting were excluded from survival analyses. Animals for cross-sectional studies were euthanized with CO2. Spleens were harvested and stored in cold PBS until analysis, and liver samples were immediately frozen in liquid nitrogen and stored at -80°C. All experiments using mice were performed in accordance with institutional guidelines for the use of laboratory animals and were approved by the Brigham and Women’s Hospital and Harvard Medical School Institutional Animal Care and Use Committees.

#### Food consumption and milk collection

To calculate food consumption by pups, dams were separated from 7-day-old pups for 6 hours; pups were weighted and returned to the dams for 1.5 hours and weighed again. Body weight change of pups’ post re-feeding was considered as a proxy for food consumption. Milk was collected from the dams when the pups were 9 days old and 14 days old. A custom mouse milking machine was constructed based on the Electric Breast Pump (Toogel). Dams were anesthetized with 3.5% isoflurane, placed on the heating pad, and injected i.p. with 100 ul of 2IU Oxytocin (Sigma) in sterile saline to stimulate milk production. 50-100 ul of milk was then collected into microtubes and frozen at -80°C. Protein concentration was measured by Qubit with a Qubit Protein Assay Kit.

#### RNA sequencing

Total RNA was extracted from snap-frozen livers using the Direct-zol RNA Miniprep Kit (Zimo) following manufacturer’s instructions. RNA was eluted with 50 ul of RNAse-free water. RNA concentration was measured with Qubit using an RNA HS Assay kit. Libraries were prepared with TruSeq Stranded mRNA LT Sample Prep Kit according to TruSeq Stranded mRNA Sample Preparation Guide, Part # 15031047 Rev. E. Libraries were quantified using the Bioanalyzer (Agilent) and sequenced with Illumina NovaSeq6000 S4 (2×150bp) (reads trimmed to 2×100bp) to get 20M read depth coverage per sample. The BCL (binary base calls) files were converted into FASTQ using the Illumina package bcl2fastq. Fastq files were mapped to the mm10 (GRCm38.p6) mouse genome, and gene counts were obtained with STAR v2.7.2b (42). Statistical analyses for gene expression were performed with custom models from *DEseq2* 3.13 and *edgeR* 3.34.1 in R (resulting lists of DEGs are in Table S3) (43). For GSEA, first ranks for each gene were calculated as -log10(p-value) multiplied by 1 if log-fold change is positive, and by -1 if negative. GSEA was performed with the *clusterprofiler* package in R (44). For signature association analysis and identification of longevity-associated genes perturbed by EL rapamycin, we filtered out genes with low number of reads, keeping only the genes with at least 10 reads in at least 50% of the samples, which resulted in 12,374 detected genes according to Entrez annotation.

#### Frailty index

Frailty index was measured as described in (26); frailty index score was calculated per the original protocol with small modifications. Because rapamycin-treated mice were significantly smaller than controls, the sd_scores for body weight and temperature were calculated in relation to the mean value of younger animals for each treatment group separately. As such, we avoided bias in these parameters due to body size differences between the groups.

#### Gait speed

Mice were positioned in the regular mouse cage that had free space on one end and a food container on the other end. Recording started with the SprinTimer app (App Store), and upon the ‘‘GO” signal from the app, mice were let run voluntarily to the other end of the cage. In the vast majority of cases, mice run towards the food container, presumably to hide. The finish line was marked by the colored tape at the end of the cage. When mice reached the finish line, recording was stopped and the time to finish was quantified based on the pictures from the app. The time of finish was determined as the time when the mouse nose crossed the finish line. Each mouse was let run four times and the median value was recorded. An attempt was considered successful if the mouse crossed the finish line without standing or turning 90 degrees or more during the run. Fifteen seconds were recorded If the mice failed to finish the run in 15 seconds.

#### Glucose and insulin tolerance tests

For the glucose tolerance test, mice were placed into food-free cages for 16 hours (usually from 7 pm until 11 am next day). The next day, mice were weighted, marked, bled from the tail and the fasting blood glucose level was measured with a glucometer. Filtered 10% glucose in sterile saline solution (Sigma) was injected i.p. at a final dose of 1 mg/kg of body weight. Injections were done at 30-second intervals between the mice. Glucose level was measured again at 20, 40, 60 and 120 minutes after the injection with a 30-second interval between the mice. Insulin tolerance test was done similarly, but mice fasted for 6 hours (usually from 10 am to 4 pm same day), and were injected with 0.75 U/kg insulin (Eli Lilly) in sterile saline solution (Sigma).

#### DNA methylation

We analyzed DNA samples isolated from the livers with the recently developed mammalian methylation array (HorvathMammalMethylChip40) (45). Two clocks were applied in our analysis, including one clock based on the mouse liver (Liver) and the other being a pan-tissue clock (Pan) (46). Student two-tailed t-test was used for statistical testing. For GSEA, first ranks for each CpG site were calculated as -log10(p-value) multiplied by 1 if log-fold change is positive, and by -1 if negative. Then ranks were aggregated by gene name, and the mean rank value for each gene was calculated. Only genes that were included in differential expression analysis were included in DNAm GSEA for comparative purposes. For GO and KEGG enrichment, genes with the sites significantly changed by treatment (p.value<0.05) were overlapped with differentially expressed genes (p.value<0.05), and an enrichment analysis of overlapped genes was carried out using all genes detected with the microarray as a background using enrichGO and enrichKEGG functions of *clusterprofiler* package.

#### Daphnia survival

*Daphnia magna* animals were used in the Daphnia lifespan assay. An IL-MI-8 heat-tolerant clone was obtained from the Ebert lab at the University of Basel, Switzerland, stock collection originating from a pond in Jerusalem, Israel. To collect synchronized cohorts, neonates born within 1–2 days were separated from the mothers, and their sex was determined at days 8–10. All mothers were also well-fed and cultured under the same conditions (25°C). We used females for all experiments in this study. After drug treatment, 40–50 animals were randomly assigned to one of our developed culture tanks (47). All cultures (e.g., mother, neonates and tank cultures) were maintained in ADaM water (48) in the 25°C incubator, exposed to a light cycle of 16 light hours followed by 8 dark hours, and fed every day the suspension of the green alga, *Scenedesmus obliquus*, at a concentration of 10^5^ cells/ml (for 1 animal/20ml density, amount of food prorated by population). Every sixth day the water was changed and the offspring were removed manually until animals were transferred to the culture platform. Details of the culture platform operation protocols are similar to those in the previously published protocols (47).

## Data availability

Raw sequencing and DNA methylation data will be deposited to GEO.

## Acknowledgments

The authors thank members of the Gladyshev laboratory for their discussion. Supported by NIH grants to VNG.

## Author Contributions

AVS designed and performed mouse experiments, interpreted the data, analyzed the data and drafted the manuscript. YC performed Daphnia experiments supervised by LP. AVS and AK conceived the project. AK, JPC, AT, JP, JH, LP and SH performed experiments, analyzed the data and revised the manuscript. VNG supervised the overall project, designed experiments, provided research materials, and wrote the manuscript.

---

**Supplementary Figure 1.**
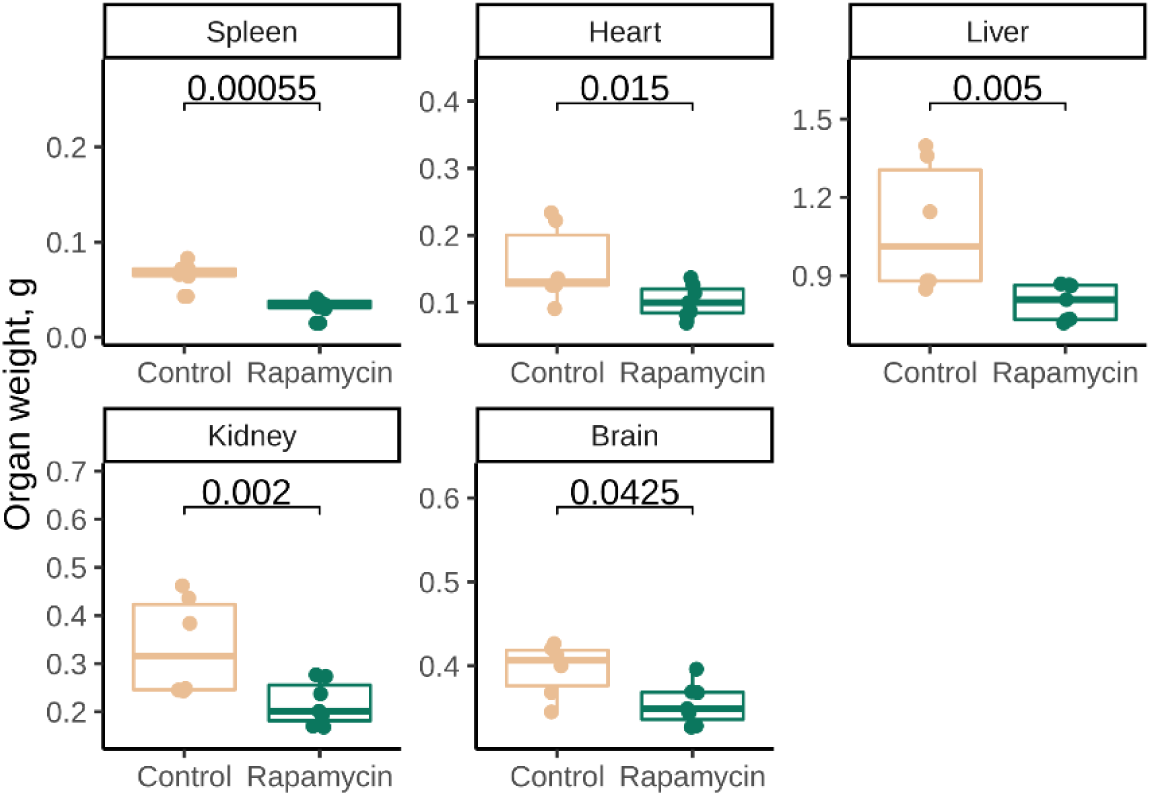
Weight of organs of 51-day-old male and female UMHET3 mice subjected to rapamycin or control diets from birth to 45 days of age. P-values are calculated with ANCOVA with sex as a covariate and are FDR-adjusted for multiple testing.

**Supplementary Figure 2.**
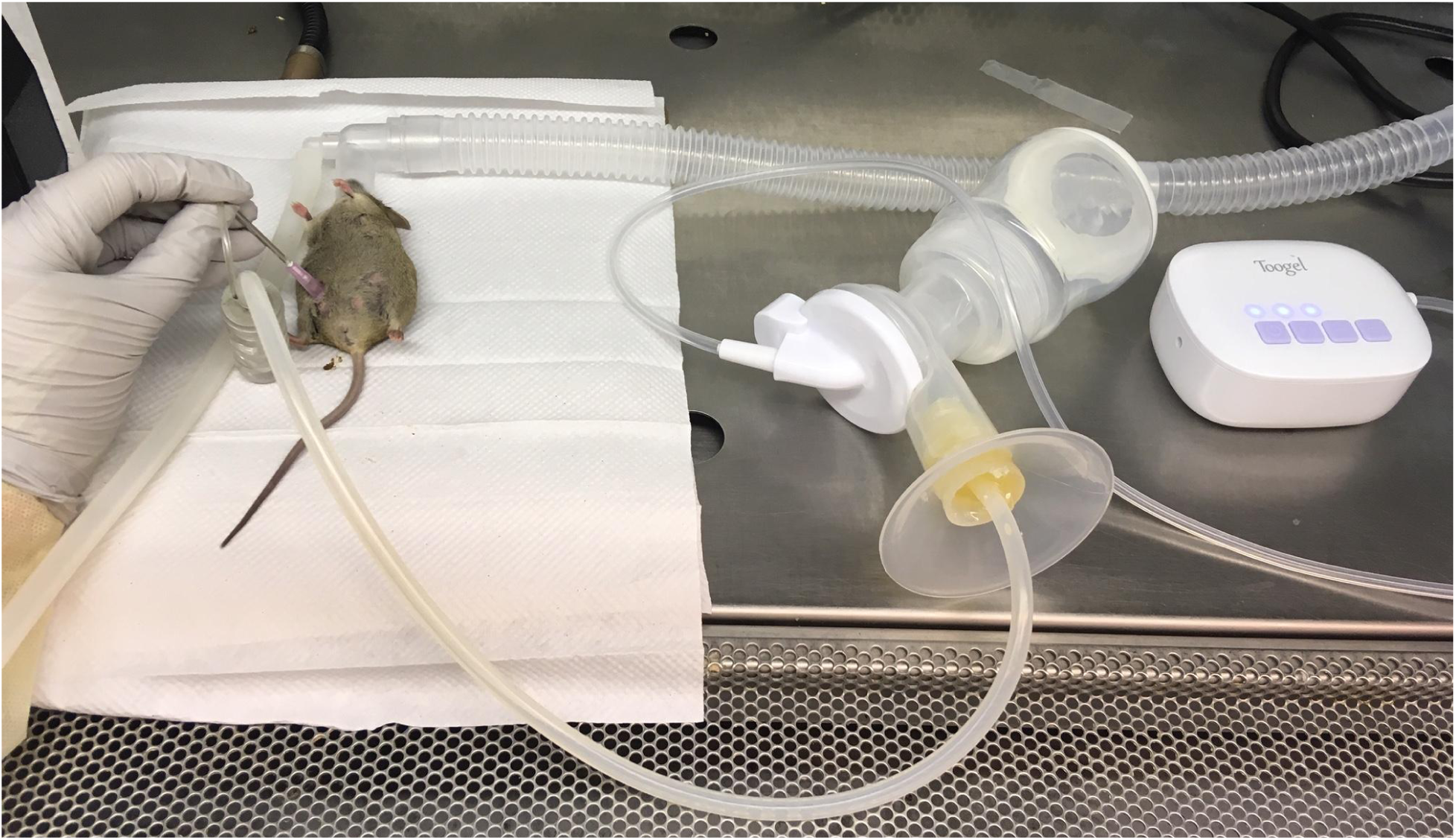
Photo of the custom setup for milk collection from mice.

**Supplementary Figure 3.**
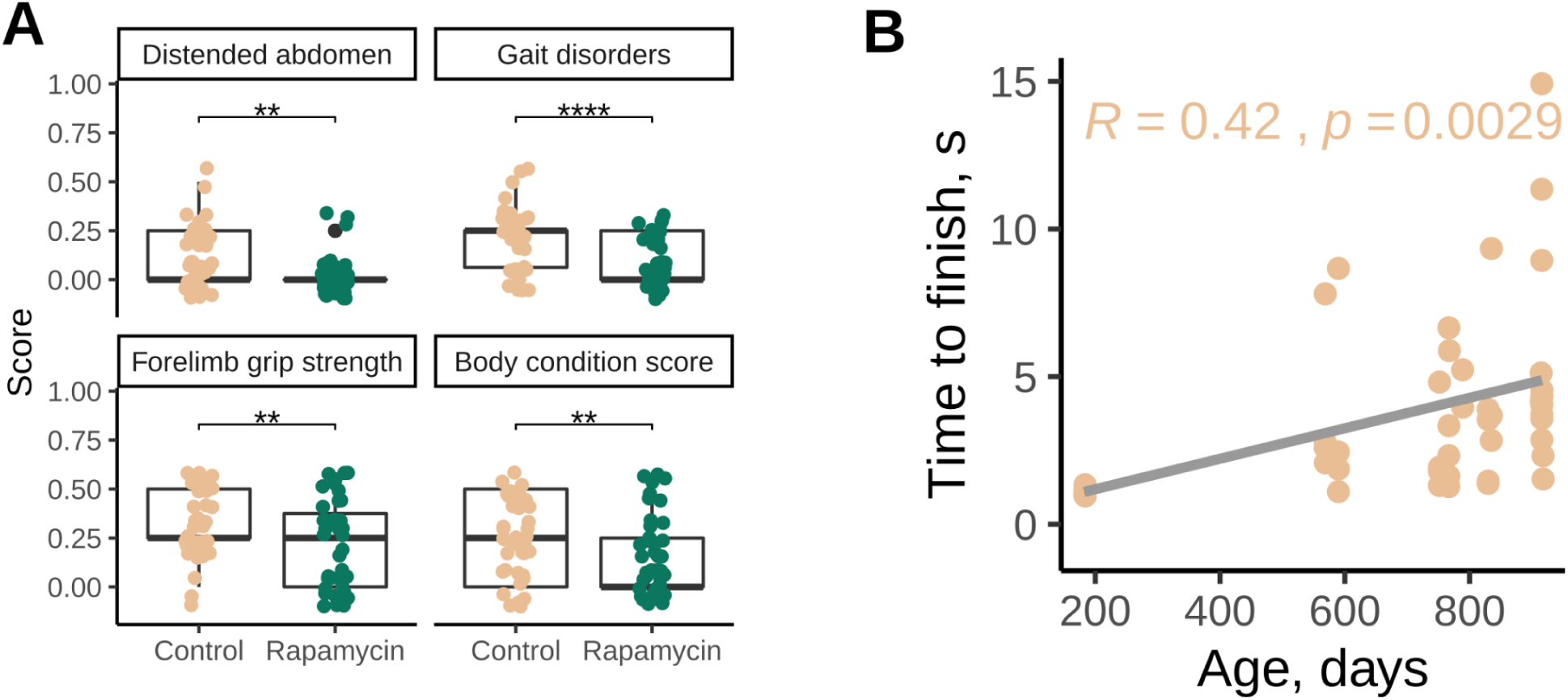
(A) Frailty index features significantly affected by rapamycin treatment. P-value is calculated with ANCOVA test with age and sex as covariates. (B) Correlation between median time to finish a run and age in UMHET3 mice.

**Supplementary Figure 4.**
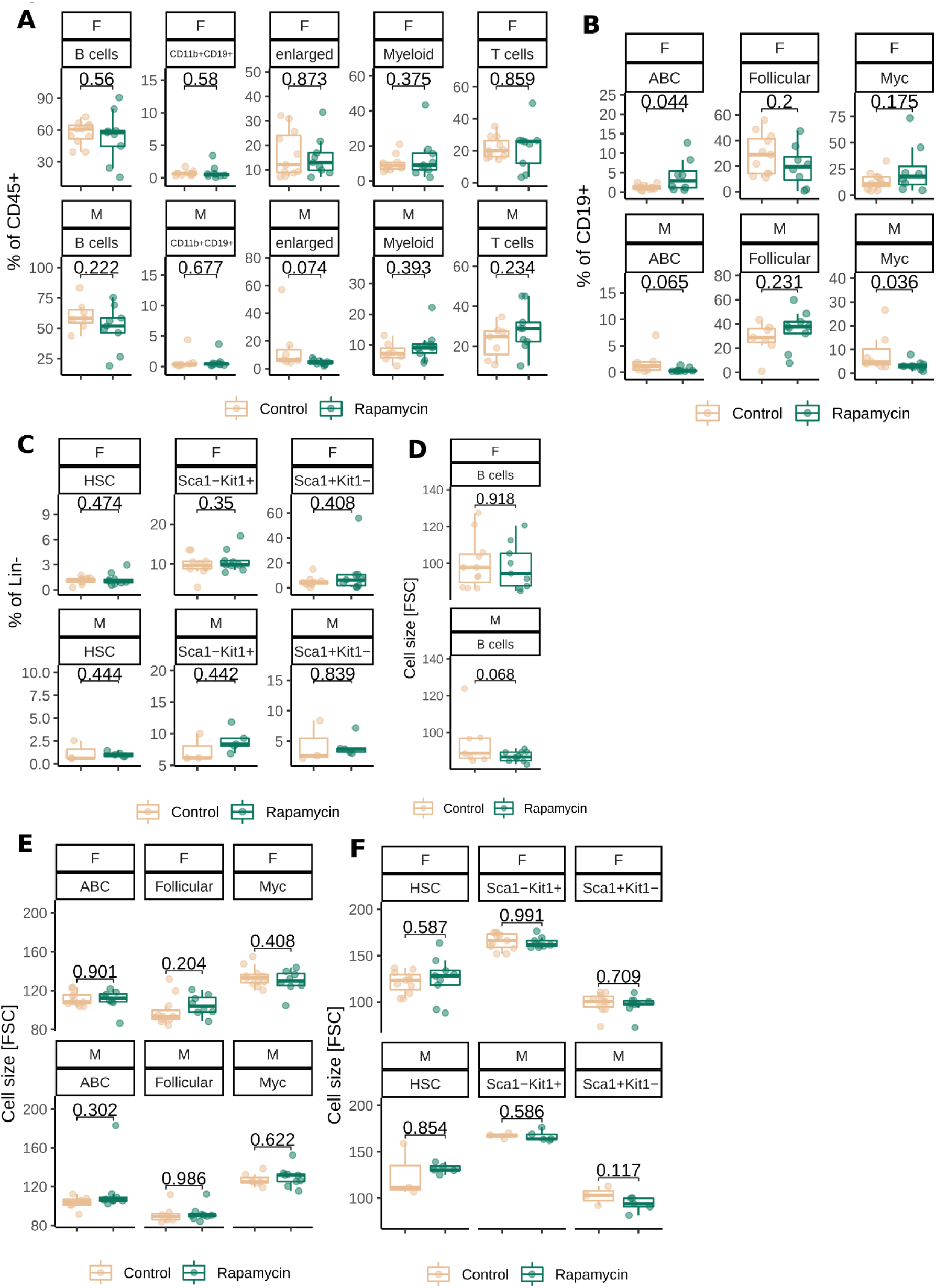
Immune cell composition and cell sizes of 21-29-day-old UMHET3 mice. P-values are calculated with ANCOVA test with age and sex as covariates.

**Supplementary Figure 5.**
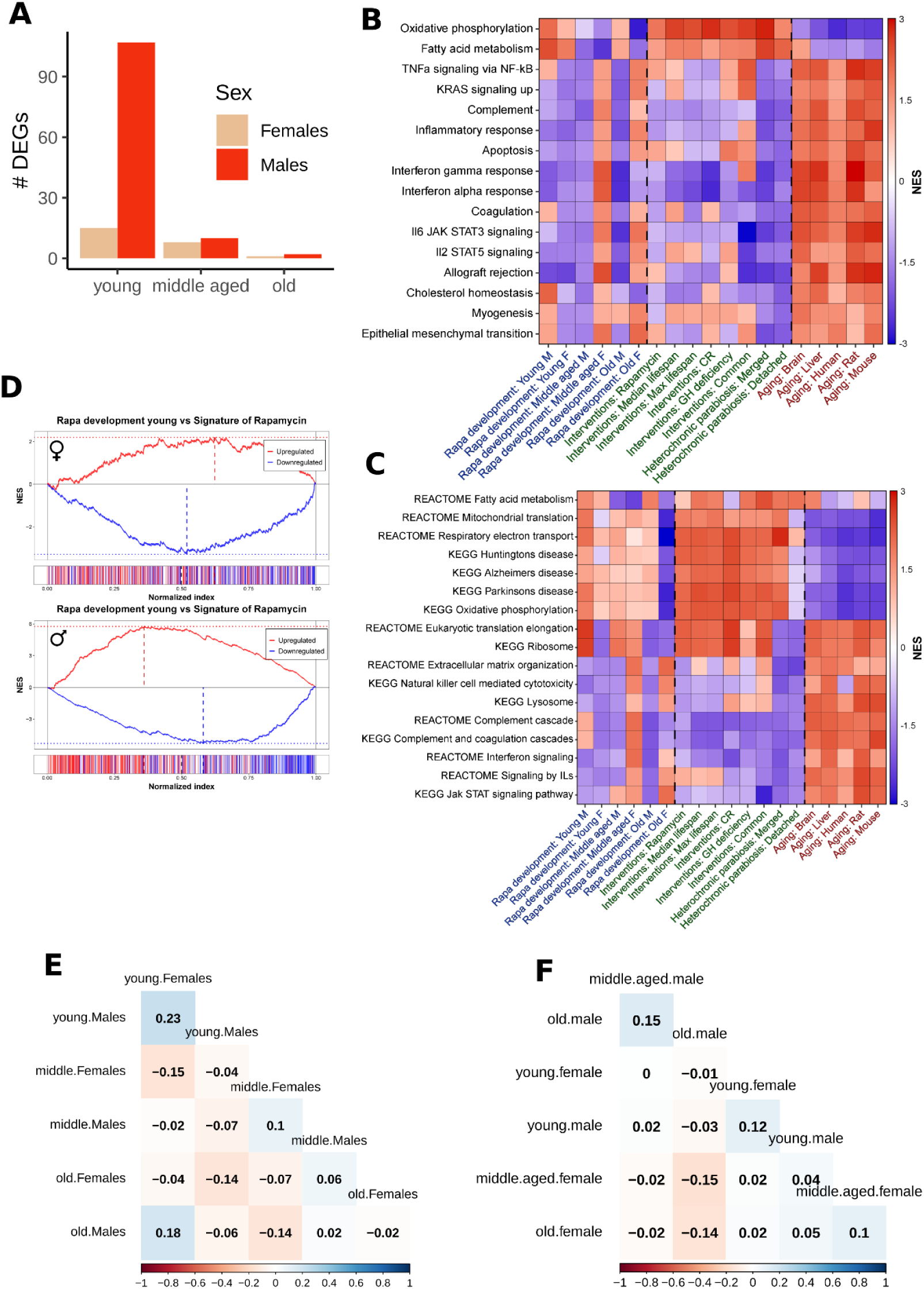
(A) Number of differentially expressed genes (DEGs, defined as FDR-adjusted p < 0.1). (B) and (C) Functional enrichment analyses of gene expression signatures and EL rapamycin treatment across age groups and sexes. Only functions significantly associated with at least one signature (q-value < 0.1) are shown. Cells are colored based on the normalized enrichment score (NES). (D) Gene set enrichment plots of young males and females against the signature of rapamycin. (E) Correlation matrix between ranks for gene expression and (F) CpG sites. Numbers displayed are spearman correlation coefficients.

**Supplementary Figure 6.**
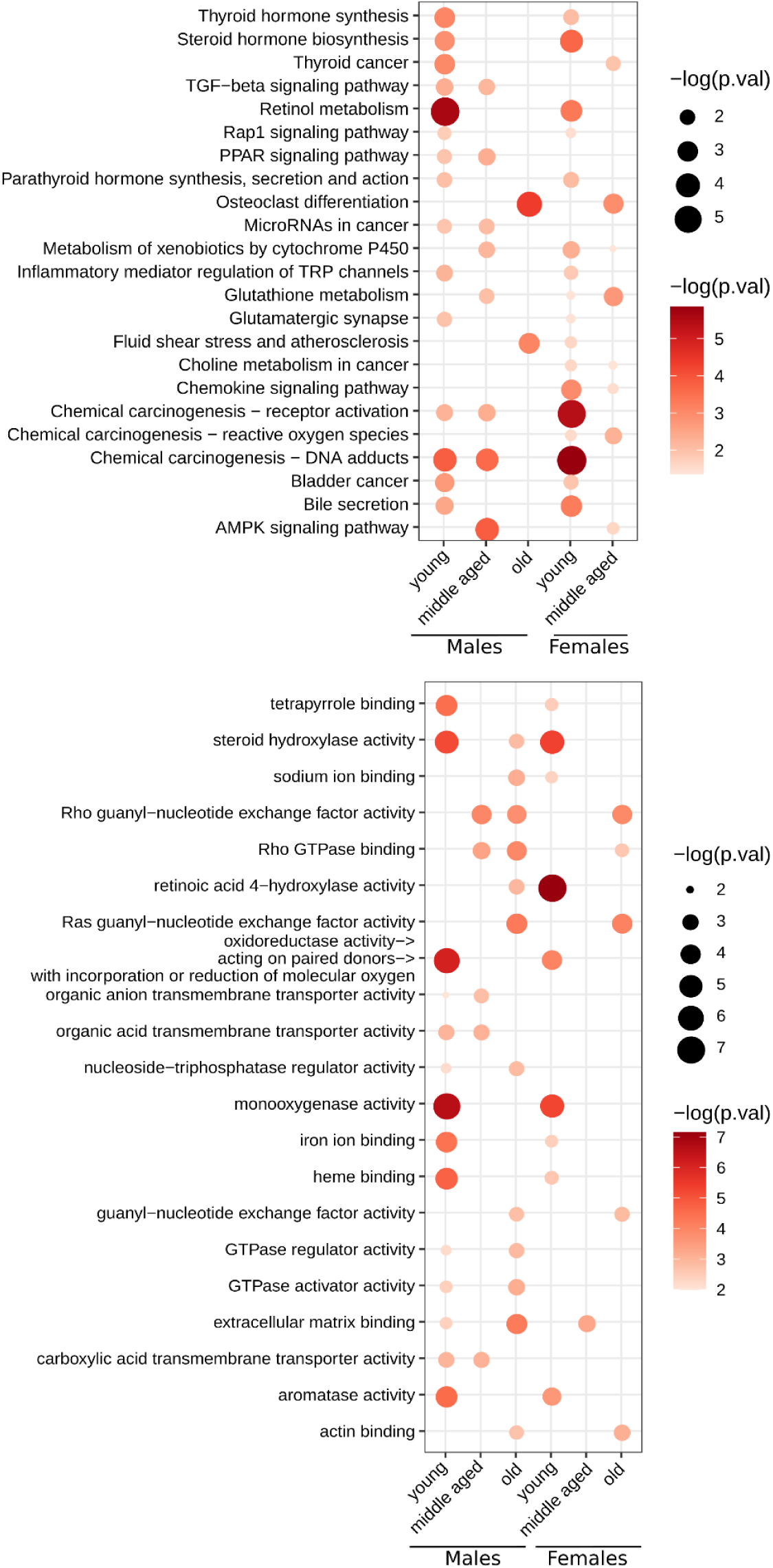
KEGG and GO enrichment of genes affected in both gene expression and DNA methylation in treated animals compared to controls. Such genes from treated old females were not enriched in any of KEGG terms.

